# Pathological beige remodeling induced by cancer cachexia depends on the disease severity and involves mainly the trans-differentiation of mature white adipocytes

**DOI:** 10.1101/2023.09.18.558327

**Authors:** Kaltinaitis B N H Santos, Pamela Knobl, Felipe Henriques, Magno A. Lopes, Felipe O. Franco, Luana L. Bueno, Stephen R. Farmer, Miguel L. Batista

**Affiliations:** Department of Integrated Biotechnology, University of Mogi das Cruzes, São Paulo, Brazil; Program in Molecular Medicine, University of Massachusetts Medical School, Worcester, Massachusetts,USA; Department of Biochemistry, School of Medicine, Boston University, Boston, MA 02215, USA

**Keywords:** cancer cachexia, adipose tissue remodeling, browning, transdifferentiation, pre-clinical models

## Abstract

In cancer associated cachexia (CAC), white adipose tissue undergoes morphofunctional and inflammatory changes that lead to tissue dysfunction and remodeling. In addition to metabolic changes in white adipose tissues (WAT), adipose tissue atrophy has been implicated in several clinical complications and poor prognoses associated with cachexia. Adipocyte atrophy may be associated with increased beige remodeling in human CAC as evidenced by the “beige remodeling” observed in preclinical models of CAC. Even though beige remodeling is associated with CAC-induced WAT dysfunction, there are still some open questions regarding their cellular origins. In this study, we investigated the development of beige remodeling in CAC from a broader perspective. In addition, we used a grading system to identify the scAT as being affected by mice weight loss early and intensely. Using different in vitro and ex-vivo techniques, we demonstrated that Lewis LLC1 cells can induce a switch from white to beige adipocytes, which is specific to this type of tumor cell. During the more advanced stages of CAC, beige adipocytes are mainly formed from the transdifferentiation of cells. According to our results, humanizing the CAC classification system is an efficient approach to defining the onset of the syndrome in a more homogeneous manner. Pathological beige remodeling occurred early in the disease course and exhibited phenotypic characteristics specific to LLC cells’ secretomes. Developing therapeutic strategies that recruit beige adipocytes in vivo may be better guided by an understanding of the cellular origins of beige adipocytes emitted by CAC.

## Introduction

Cancer associated cachexia (CAC) is characterized by body weight loss, systemic inflammation, remodeling of adipose tissue (AT), and skeletal muscle wasting [1]. This affects the quality of life, the survival rate, and the complications associated with cancer treatment. Around 20% to 30% of cancer patients die from cachexia [2]. AT metabolism is altered in CAC, resulting in low fat mass and poor survival rates [3, 4]. Furthermore, a clinical longitudinal study had demonstrated that in patients with progressive cancer cachexia, body fat was lost more rapidly than lean tissue, highlighting the importance of considering AT dysfunction when planning effective palliative care [5]. In CAC, the white adipose tissue (WAT) undergoes morphofunctional and inflammatory changes, which result in tissue dysfunction and remodeling [1, 6-9]. Studies using different experimental models and a longitudinal approach have characterized the development process of WAT remodeling, which have demonstrated that the dysfunction of genes expression and adipokines involved in normal WAT function are already present before any morphological changes detection, for example, adipocyte atrophy [2, 10]. There is also consistent evidence that this phenotype is present both in clinical studies [2, 6, 11, 12] and in multi-experimental CAC models [8, 13].

It has recently been demonstrated that “niches’’ of beige adipocytes (UCP1+ multilocular cells) can be found within WAT in association with the changes described above that characterize WAT remodeling. Aside from this phenotypic picture, an increase in WAT thermogenesis has been identified in the literature as an important contributor to accelerated energy expenditure and weight loss in CAC mouse models [8, 9]. The “beige remodeling” occurs in preclinical models of CAC when WAT cells gradually differentiate into Brown Adipose Tissue (BAT)-like cells, also referred to as “beige” cells [1, 8, 9]. CAC has been shown to switch white adipocytes in WAT from mice and humans [3, 9] to beige adipocytes. The observation of increased UCP1+ multilocular cells from CAC indicates that adipocyte atrophy may be associated with increased beige remodeling in human cancer cachexia. Additionally, considering the relevance of beige remodeling as a relevant event in the different stages of cachexia, pharmacological treatment with β-adrenergic antagonists, nonsteroidal antiinflammatory drugs (Sunlidac), and TLR4 antagonists (Simvastatin) has been shown to effectively reduce beige remodeling in CAC in tumor-bearing animals while preserving body mass [1]. Only the latter showed significant effects and increased survival. However, in spite of the relevance of beige remodeling in CAC-induced WAT dysfunction, a number of open questions remain regarding the cellular origins of beige adipocytes and whether this is related to tumor type or experimental model. Developing therapeutic strategies that recruit beige adipocytes in vivo may be better guided by a detailed understanding of the cellular origins of beige adipocytes emitted in response to CAC stimuli.

In this study, we sought to identify and characterize more broadly the development of beige remodeling in CAC. The early stages of beige remodeling in WAT were specific to the anatomical region of subcutaneous adipose tissue. The phenotype was only evident in animals that had developed a substantial amount of CAC. Furthermore, we have demonstrated using different in vitro and ex-vivo techniques that the secretome of Lewis LLC1 cells can induce a switch from white to beige adipocytes, which is specific to this type of tumor cell. These beige adipocytes are predominantly the result of cell transdifferentiation during the more advanced stages of CAC.

## Methods

### Animal model

Eight-week-old male C56BL/6 (Jackson laboratory # 000664), (wild type, weighing approximately 20-25 grams were used. The animals were obtained from The Jackson Laboratory (Maine, United States of America), and maintained by the central vivarium of the University of Mogi das Cruzes (Ceua # 010/2019 / # 006/2017 / 016/2016 and 01/2017). The animals were kept in individual cages, at an average temperature of 22°C ± 1ºC, or in a cold chamber, at 4°C ± 1ºC, for 7 days (prolonged exposure to cold), with light/dark cycles every 12 hours, in a controlled manner, besides the provision of feed (Nuvilab®, Nuvital S/A, Colombo, PR) and water ad libitum. UCP1-Cre mice and B6.129(Cg)-Gt(ROSA)26Sortm4(ACTB-tdTomato,-EGFP)Luo/J (mTmG) mice are from Jackson Laboratory. The Ucp1-cre mice were crossed with mTmG mice to remove the flox-flanked resistance cassette. Mice were housed at 23°C in a 12 hr light/dark cycle with free access to normal chow. All experiments used matched littermates. Experimental procedures were approved by the Boston University Institutional Animal Care and Use Committee.

### Tumor cells inoculation

Lewis Lung Carcinoma (LLC1) tumor cells (LLC1, CRL-1642 -ATCC) were used for inducing cancer cachexia syndrome. LLC1 was injected subcutaneously into the right flanks (200 μl LLC cells −3.5 × 10^5^). Non-tumor bearing control mice received Phosphate-buffered saline (PBS) only. Development of cachexia was monitored by tumor size and body weight for 28 days [2,14, 15].

### In Vitro models

Swiss preadipocytes 3T3-L1 cell line (ATCC CL173 -American Type Culture Collection, Manassas, VA, USA), were plated at 1x10^4^ in 24-well culture plates and differentiated according to [43]. Stromal vascular fraction (SVF) cells and adipocyte isolation were adapted from Neves and coworkers (Neves *et al*. 2015), as well as Rodbell’s method (Rodbell 1964) with minor modifications (see supplementary methods). Freshly isolated mature adipocytes from UCP1-Cre mice were cultivated for 48 hours using 6.5mm Transwells (Costar-3413 and 3397) with a minor adaptation.

### Tumor cells and Conditioned medium preparation

Lung Lewis carcinoma (LLC1, CRL-1642 -ATCC), colorectal carcinoma 26 (C26, CT26.WT - CRL-2638 -ATCC) and colon adenocarcinoma (MC-38, CVCL_B288) tumor cells, were cultured in Dulbecco’s Modified Eagle Medium (DMEM), (Gibco,11965084), with 4500 mg glucose/L, L-glutamine (4 mM), NaHCO3 (2mM), supplemented with 10% bovine serum (Gibco, 16170) and 1% penicillin and streptomycin (Gibco, 15140122), at pH 7.4 and maintained in an oven at 37°C with 5% CO_2_.For conditioned medium collection, cells were plated at a 1:3 confluence. The following day the medium was changed to the Freestyle expression medium (Invitrogen) and was then collected 24 h later. For fractionation, the Tumor-cell-conditioned medium was filtered using a 3-kDa cut-off filter (Millipore). Freshly adipocytes were treated with conditioned medium for 48 h after cell isolation. SVF started at day 0 of the differentiation protocol. When cells were treated with conditioned medium, the treatment medium was composed of 50% fresh adipocyte culture medium and 50% LLC-cell-conditioned medium. When the conditioned medium was filtered and concentrated, adipocytes were exposed to 75-50% adipocyte/SVF culture medium and 25-50% concentrated tumor-cell conditioned medium.

### Mice weight loss grading system

After the inoculation of tumor cells, the animals were kept for an experimental period of 28 days, the time required for the development of the syndrome [17]. The evolution of the syndrome was monitored through the individual measurement of the animals’ body weight (BW) (OHAUS® USA). Initially, in order to classify the animals according to the degree of cachexia, according to the magnitude of the BW loss, we used a database composed of C57BL/6 animals; 26 controls (saline-inoculated) and 25 animals inoculated with 3.5 x 10^5^ LLC1 cells, for 28 days. The classification of the magnitude of the development of the cachexia syndrome was established by subtracting the measured value of BW on day 28 by the highest value of body mass measured during the experimental period. For plateau detection, at least two measurements with reduced and subsequent values should be detected. Thus, the experimental groups were classified into; Control (CO), being the animals inoculated with saline solution; Pre Cachexia (PC), with a reduction of up to 5% of body mass, Cachexia (CC), defined with a reduction of ≥5% to 20% of body mass, Severe Cachexia (S-CC), which present a reduction ≥20% of body mass, at the end of the experiments. For the cachexia index, the equation described by [18] was used.

### Statistical analysis

Data was analyzed in GraphPad Prism 8 (GraphPad Software). The statistical significance of the differences in the means of experimental groups was determined by Student’s t-test or and 2-way ANOVA, followed by Tukey’s post hoc comparison tests. The first observation indicates that the results presented a normal distribution, after performing the Kolmogorov-Smirnov test, and that there is no difference in variance between the group samples, after performing the equality of variance test. Pearson’s coefficient of variation (r) was used as the average of the degree of linear relationship between the quantitative variables. The data are presented as means ± SEM. P values ≤ 0.05 were considered significant.

## RESULTS

### Characterization of cancer cachexia using “mice weight loss grading system”

The profile of CAC development was characterized using data obtained from 51 animals collected between 2012 and 2016 [2]. To begin the classification of cachexia staging, descriptive analysis and normality distribution regarding CAC features were conducted (Table S1). The first analysis verified an increase in BW (somatic growth). At the end of the experimental period (day 28), BW gain was significantly lower in the CC group (6 %) than in the Control group (CO). (S1 A). Furthermore, the CC group presents a shift to the left of normal distribution in terms of BW gain average (S1 B). Only a few animals from CC showed BW values lower than those measured at the initial day of the experiment. As a result of this scenario, it is evident that it is necessary to correct the “body weight effect due to somatic growth” factor in tumor-bearing individuals. This was normalized by adding the higher BW values from the somatic growth plateau (plateau) as a relativizing factor (S1 C). After correction, 35% of body BW loss was observed in the CC group (S1 D). Using this new classification system, it is easy to distinguish between animals that develop CC and those that did not (S1 E). There were no tumor mass differences associated with the syndrome (S1 F). Table 1 shows CAC features. Most individuals were classified as CC (Cancer Cachexia), thus representing 60%, followed by the PC (Pre-Cachexia) group (24%), and finally a group of animals affected by severe-cachexia (S-CC, 16%). The following analysis was conducted using a mice weight loss grading system. In order to do so, the animals were separated based on the severity of cachexia syndrome; the control animals without tumors were CO (Controls).The experimental groups were classified as follows: Control (CO), which comprised animals that were injected with saline solution; Pre-Cachexia (PC), which resulted in a reduction of up to 5% of body mass, Cachexia (CC), which resulted in a reduction of up to 20% of body mass, Severe Cachexia (S-CC), which resulted in a reduction of up to 20% of body mass. The Voltarelli [18] equation was used to calculate cachexia index (Fig A1).

Tumor-Bearing Mice
| Parameters | Control |  |  | Pre Cachexia |  |  | Cancer cachexia |  |  | Severe cancer cachexia |  |  | Tukey |
| --- | --- | --- | --- | --- | --- | --- | --- | --- | --- | --- | --- | --- | --- |
|  | Mean | Dp | N | Mean | Dp | N | Mean | Dp | N | Mean | Dp | N |  |
| Plateau BW (Day) | 28.0 | 0.0 | 26 | 27.5 | 0.5 | 6 | 25.4 | 4.5 | 15 | 23.6 | 4.0 | 4 | Cc x Co (p=0.020); S-CC x Co (p=0.017) |
| Tumor Apperance (Day) | - | - | - | 13.3 | 0.8 | 6 | 13.9 | 1.9 | 15 | 13.2 | 2.3 | 4 | - |
| Tumor Weigth (g) | - | - | - | 0.8 | 0.3 | 6 | 1.4 | 0.5 | 15 | 2.1 | 1.0 | 4 | S-CCx PC (p=0.0045) |
| Delta (%) | 15.6 | 8.6 | 26 | 12.7 | 6.6 | 6 | 8.0 | 6.4 | 15 | 2.4 | 5.9 | 4 | Cc x Co (p=0.018); S-CC x Co (p=0.012) |
| Carcass - Plateau (g) | - | - | - | -0.9 | 0.2 | 6 | -2.4 | 0.9 | 15 | -7.1 | 1.2 | 4 | CC x PC (p=0.0038); S-CC x PC (p= <0.0001); S-CCx CC (p= <0.0001) |
| Carcass - Plateau (%) | - | - | - | -3.7 | 0.8 | 6 | -9.7 | 3.5 | 15 | -26.2 | 4.1 | 4 | CC x PC (p=0.0022); S-CC x PC (p= <0.0001); S-CC x CC (p= <0.0001) |
| Cachexia Index | - | - | - | 1.7 | 5.6 | 6 | 6.2 | 5.4 | 15 | 11.8 | 5.0 | 4 | S-CC x PC (p=0.021) |
| Tumor x BW -1 (g) |  |  |  | 3.33 | 1.42 | 6 | 6.20 | 2.42 | 15 | 9.93 | 4.64 | 4 | S-CC x PC (p=0.0025) |
| Rectal Temperature | 37.5°C | 0.32 | 26 | 36.9°C | 0.6 | 6 | 36.5°C | 0.7 | 15 | 32.4°C | 2.7 | 4 | S-CC x CO (0.0231) |

BW, Body Weight;

BW loss is the main clinical marker of cachexia [22-25]. When compared to CO, consistently carcass and BW losses were showed in the CC and S-CC (12% and 45% respectively) (*Figure* 1A, B). *Figure* 1 (C-E) shows the general readouts of the mice weight loss grading system. BW plateau was used as a normalizing factor to compare all groups to CO (*Figure* 1C). The tumor mass in the S-CC group is 50% greater than that in the CC group, which also suffers from the greatest loss of BW. As compared to the PC group, both the CC and S-CC groups (52% and 89% increase when compared to PC) presented a high cachexia index (*Figure* 1E). Furthermore, white adipose tissue (WAT) has consistently been reported as an early affected by CAC [6, 11, 26]. *Figure* 1 (F-I) shows different WAT depots; subcutaneous (scAT), visceral fat pads (rpAT and epAT) in different cachexia stages. In comparison with CO animals, CAC caused wasting of fat pads in S-CC (scAT 21.1%, epAT 16.3%, and rpAT 10.56%). Additionally, scAT was early (CC group, 50%) and more intensely affected (in terms of tissue wasting) than visceral AT (epAT and meAT). Only the scAT shows a negative correlation (Pearson = 0.0269, p<0.05) with the BW loss.

**Figure 1.**
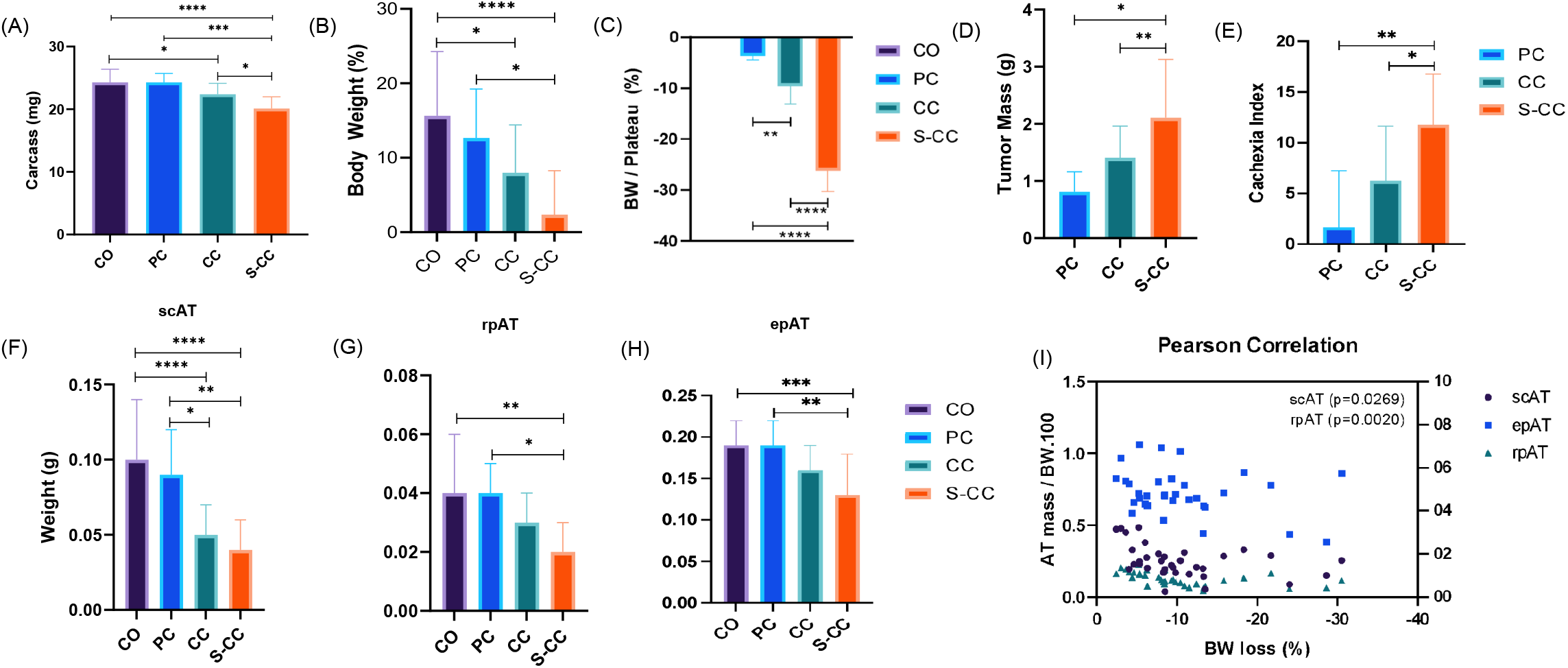

### WAT remodeling in CAC

Using a grading system for mice weight loss, we demonstrated that the scAT is affected early and intensely by mice wasting. Our next step was to examine the morphofunctional parameters of the scAT in a broader context. To accomplish this, the scAT was divided into two distinct anatomical regions, namely, the dorsolumbar region and the inguinal region [27] (*Figure* 2A). This approach revealed that the diameter of cells in CC animals was reduced only in the inguinal region (*Figure* 2B), which is followed by an increase in the percentage of adipocytes with a diameter of less than 200 μm^2^ and higher frequency of the smaller adipocytes (*Figure* 2C). In contrast, in the dorsal lumbar region, adipocytes are distributed similarly to those in the CO group (*Figure* 2D and E). The collagen content of scAT shows ECM changes in CAC. As compared to the control group, collagen deposition increased by 28% in the CC group (*Figure* 2F). *Figure* 2G shows a 16-fold increase in fibronectin expression (*Fn*) in adipocytes in the CC and S-CC groups. CC showed increased gene expression in both lipolytic and lipogenic processes of *Atgl, Lipe, Plin, Adbr3, Glut4, Fas, Pck1*. Despite this, the *Acsl1* gene (long-chain fatty acid -CoA ligase 1), which plays a key role in lipid biosynthesis, and fatty acid degradation, was down-regulated in S-CC animals. Additionally, the inflammatory profile of the scAT was evaluated. All tumor-bearing groups (PC, CC, S-CC) have CD68 (macrophages) positive cells in both anatomical regions. Further, inflammatory gene profiles in scAT adipocytes and SVF were assessed (*Figure* S2 E). Inflammatory gene markers (TNF, IL-10 and ILb) expression increase as the syndrome progresses in all 3 experimental groups.

**Figure 2.**
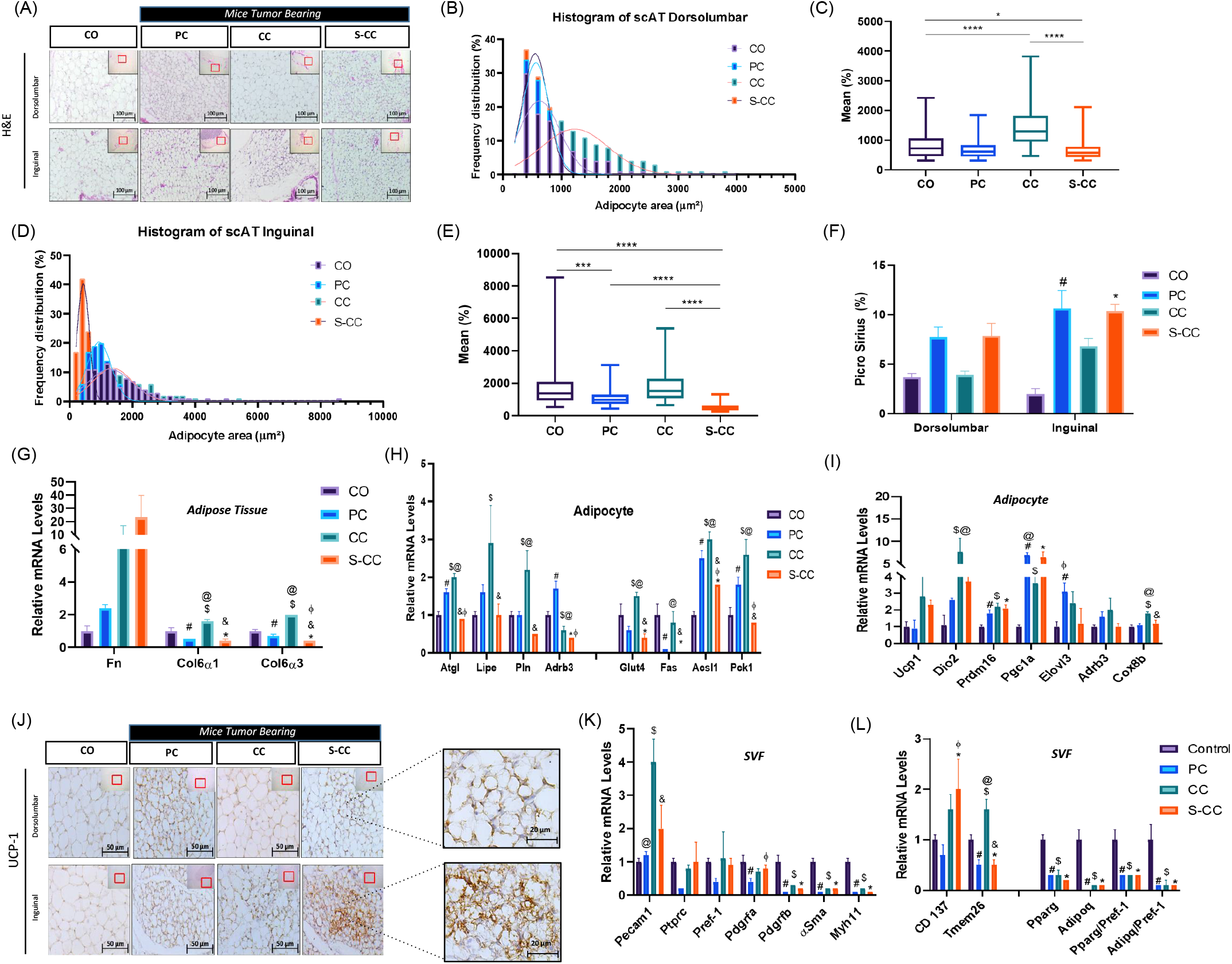

### Pathological beige remodeling induced by CAC

To compare the browning phenotype of the WAT in CAC, we employed prolonged cold exposure as a benchmark (*Figure* S2 A). S-CC animals display BAT dysfunction and reduced rectal temperature, which correlate with BW reduction. Hence, we used the S-CC group for phenotypic and comparative analyses. The morphofunctional changes leading to WAT remodeling are well described in CAC [1, 6, 12, 28, 29]; however, beige cells (browning) appear to depend on the animal model and experimental conditions. Considering this, we compared the presence of UCP1-expressing cells in different experimental groups after scAT remodeling. In *Figure* J2, S-CC exhibits an abundance of “islands” composed of small adipocytes with large nuclei, multilocular cytoplasm, and positive staining for UCP-1. In S-CC animals, UCP-1 positive cells are predominantly increased in the inguinal region and more intense in the lumbar dorsal region. The rectal temperature was reduced in S-CC mice housed at standard temperatures (23°C) despite WAT browning increasing thermogenesis. BAT surface areas were analyzed for temperature control (directly) and heat dissipation prior to severe cachexia. *Figure* S2 F and G show that ROI temperature in the BAT region decreased by 6% (p=0.06) in animals suffering from the most severe form of the syndrome (S-CC). S-CC animals have a 13% (p=0.06) lower rectal temperature than the other experimental groups (*Figure* S2 H). Even though tumor growth continued to increase in both experimental groups, no statistical difference could be observed (*Figure* S2 I).

### De novo beige adipogenesis

Using freshly isolated mature adipocytes from scAT, we examined the main thermogenic markers profile in context of CAC (*Figure* 2K). *Prdm16, Pgc1α, and Elovl3* were increased in all experimental groups. The next step was to examine the stromal vascular fraction (SVF) cell population once beige adipocytes were detected. This analysis aimed to determine the density of progenitor cells that may be involved in differentiation into beige cells. The expression levels of *Pdgfrb* (adipogenic-fibroblastic progenitor), *αSma*, and *Myh11* (mural progenitors) were shown to be reduced as a function of CAC (PC, CC, S-CC), with no changes in *Pdgfra* (adipogenic progenitor). Similarly, *Figure* 2L indicates adipogenesis-related gene expression reductions, in this case, markers of white adipogenesis. The presence of genes (*Cd137* and *Tmem26*) used as markers of beige adipocytes commitment in the population of SVF cells was also evaluated [30-33]. These results indicate an increased progenitor cell (*Cd137*) in the S-CC group, might suggesting increased cells committed to beige adipogenesis in the SVF. In turn, mural progenitors, or those not committed to “white” adipogenesis, showed reduced levels of their respective mRNA. In this regard, it is possible to speculate that these changes in density might affect both white and beige adipocyte differentiation (*de novo*).

In recent years, CAC induces beige cells in WAT has been described [1, 2, 8, 9]. Even so, there are still several questions to be answered, including how different experimental models and tumor secretome profiles are involved. We used co-culture approach to study how Lewis LLC1 cells secrete related products might modulate adipogenesis (*Figure* 3A-D). To gain more information about LLC1 cells’ specificity, conditioned medium containing secretome from different tumor cells, such as colon carcinoma 26 (C-26) and colon adenocarcinoma MC38, was added to the assays. Post-stimulation morphology is shown in *Figure* 3E. The cell density has decreased, as expected. In this parameter, the assays performed with tumor cells in conditioned medium did not show a marked difference. A 10-fold increase in the expression of UCP-1, the main beige cell marker for thermogenesis dependent on UCP-1, was observed in W+LLC when compared to cells without conditioned medium (MID-W) or assays with C26 when evaluating thermogenic genes (*Figure* 3F) and progenitor cell markers (*Figure* 3G). In particular, the genes *Pgc1aα, Cptb, and Cox8b* were increased in W+LLC1 relative to all other experimental conditions. Although W+C26 increased slightly changes in thermogenic gene expression, the effect of CM medium with LLC was much higher than in the other tumor cells. In *Figure* 3C, we show the expression of markers used to identify adipogenic progenitor cells, murals, and fibroblasts (*Fn1, Col1α1*, and *Col6α3*). *Fdgfra* expression seems to be more sensitive to CM from MC38 and *αSma* than to cultivating with CM alone. As for fibroblast markers, *Col1α1* and *Col6α3* showed a very distinct profile, like non-differentiated cells for W+LLC and W+C26. Moreover, to the best of our knowledge, this is the first study that investigated the possible participation of progenitor (non-classical) cells in beige adipogenesis in CAC. Considering the above results, we wondered if the primary cells would differentiate only with the addition of conditioned medium with tumor cells (CM), without the addition of differentiation-inducing cocktail (adipogenesis) (*Figure* 3H). This assay used CM produced with LLC1 (CM-LLC1). Even in small numbers, primary cells differentiated and acquired mature adipocyte morphology (*Figure* 3H), followed by an increase in beige adipocyte marker gene expression (*Figure*s I3 and J).

**Figure 3.**
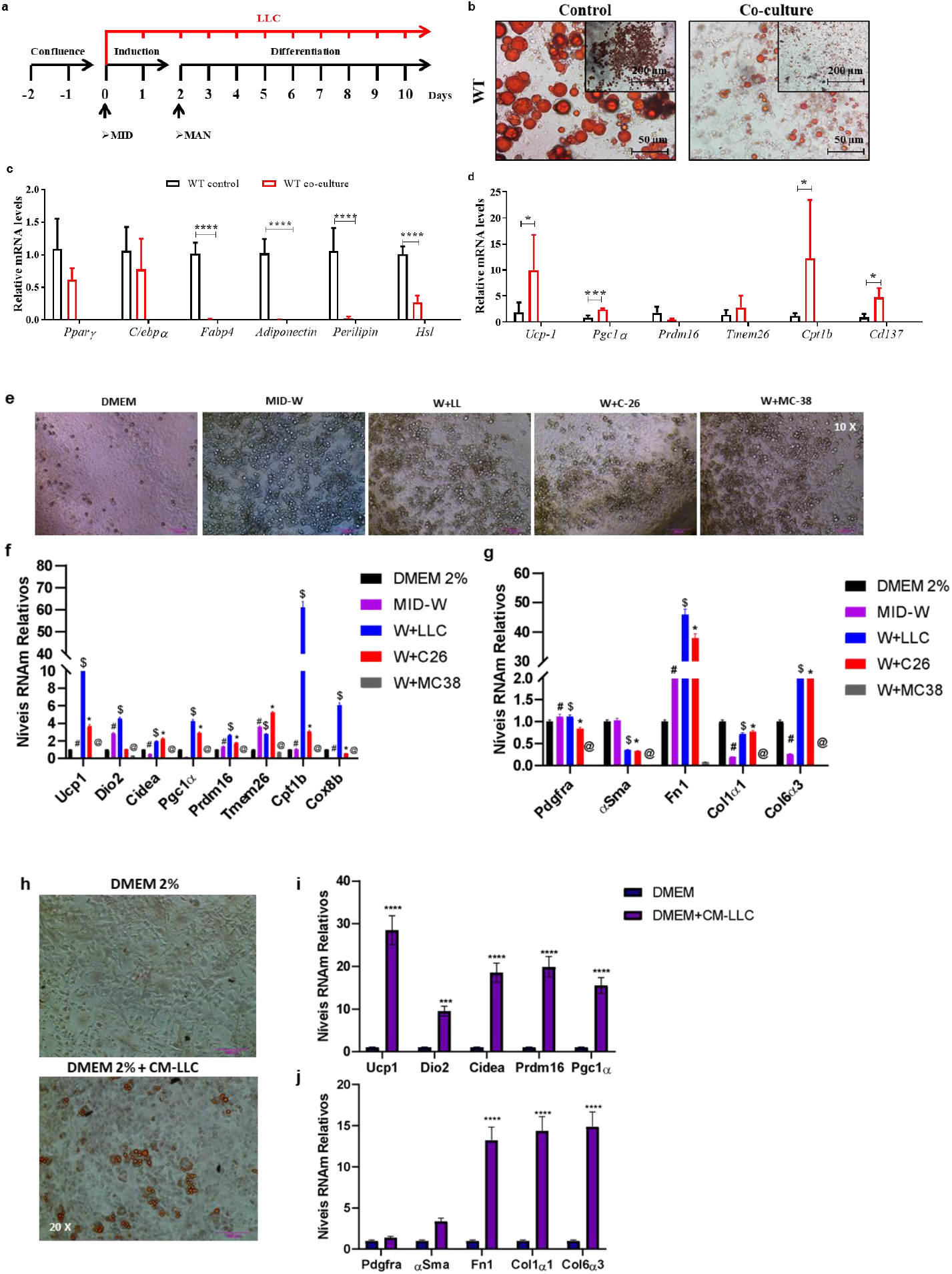

### Transdifferentiation

Aside from de novo adipogenesis analyses, trans-differentiation has also been described as a potential pathway for generating beige adipocytes [8]. In general, and not for too long, this process has been described as relevant to the browning phenotype, particularly when induced by the application of an adrenergic agonist (CL-316,243) [34, 35]. Further, given the general scenario of CAC, particularly the remodeling and dysfunction of WAT, this process could play a significant role in the development of cachexia. Our findings were further enhanced by using UCP1-Cre mice to characterize the presence of UCP-1 positive cells both in vivo and ex vivo (*Figure* 4A). We used mature adipocyte aggregate assays (MAAC) in culture with transwell plates in *Figure* 4A [36]. Therefore, in the next step, we used primary freshly isolated mature adipocytes from UCP1-Cre mice as an experimental model to study trans-differentiation in fresh mature adipocytes. Therefore, this approach would be the ideal technical procedure for determining whether LLC1 cells induce transdifferentiation of white to beige adipocytes. As shown in *Figure* 4B, the MAAC assay was cultured for 48 hours with CM-LLC1 (25%). After treatment with CM-LLC1, a greater number of GFP-positive cells (green) are observed, indicating a greater number of UCP1-GFP cells. In the control condition, no GFP+ positive cells were observed. As expected, when we increased the CM-LLC1 contraction in the assay from 25 to 50%, both the number of UCP1-GFP+ cells and the expression of thermogenic genes increased (*Figure* 4C and D). According to *Figure* 4E, CM-C26 did not produce significant changes under the same experimental conditions.

**Figure 4.**
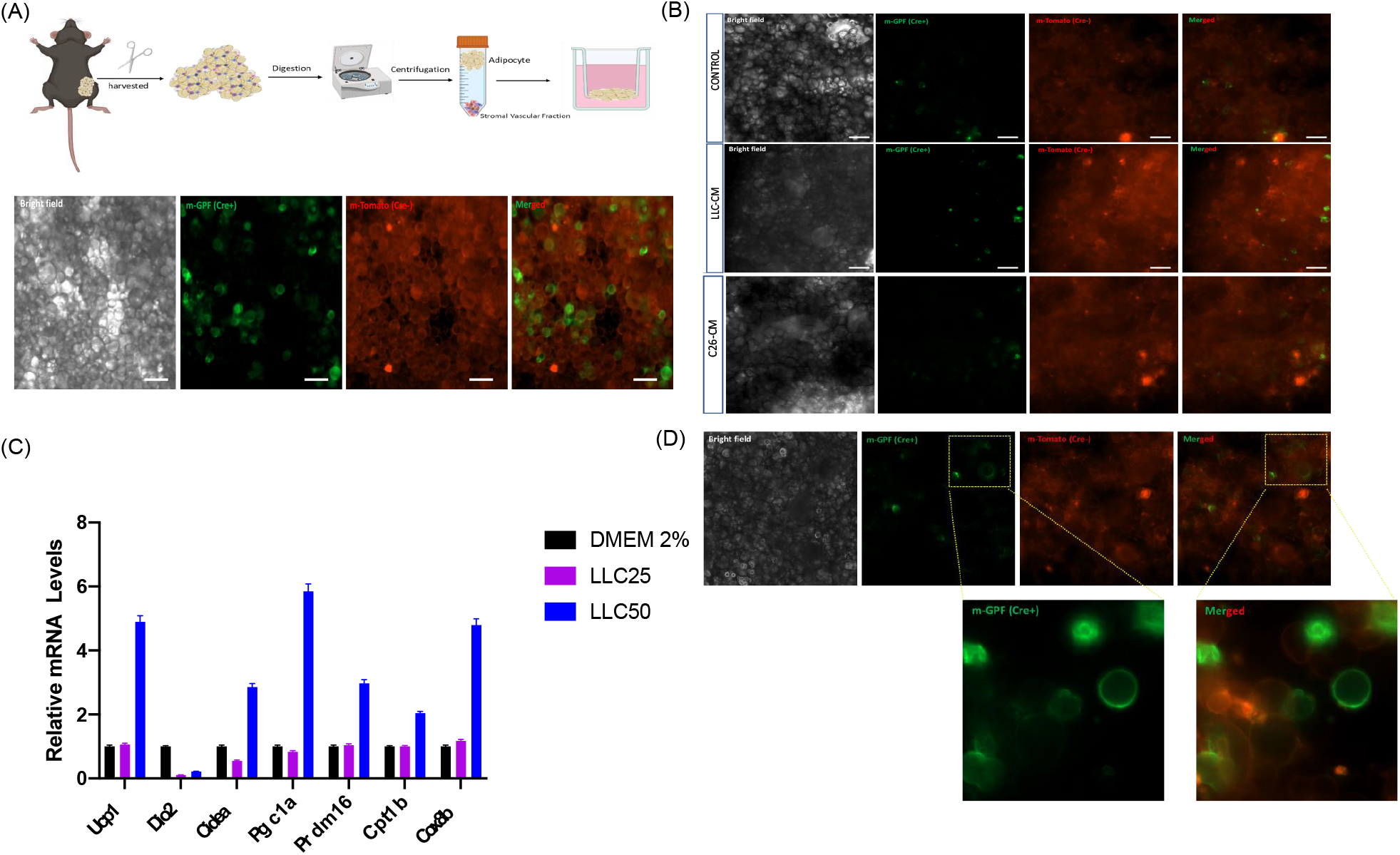

**Figure 5.**
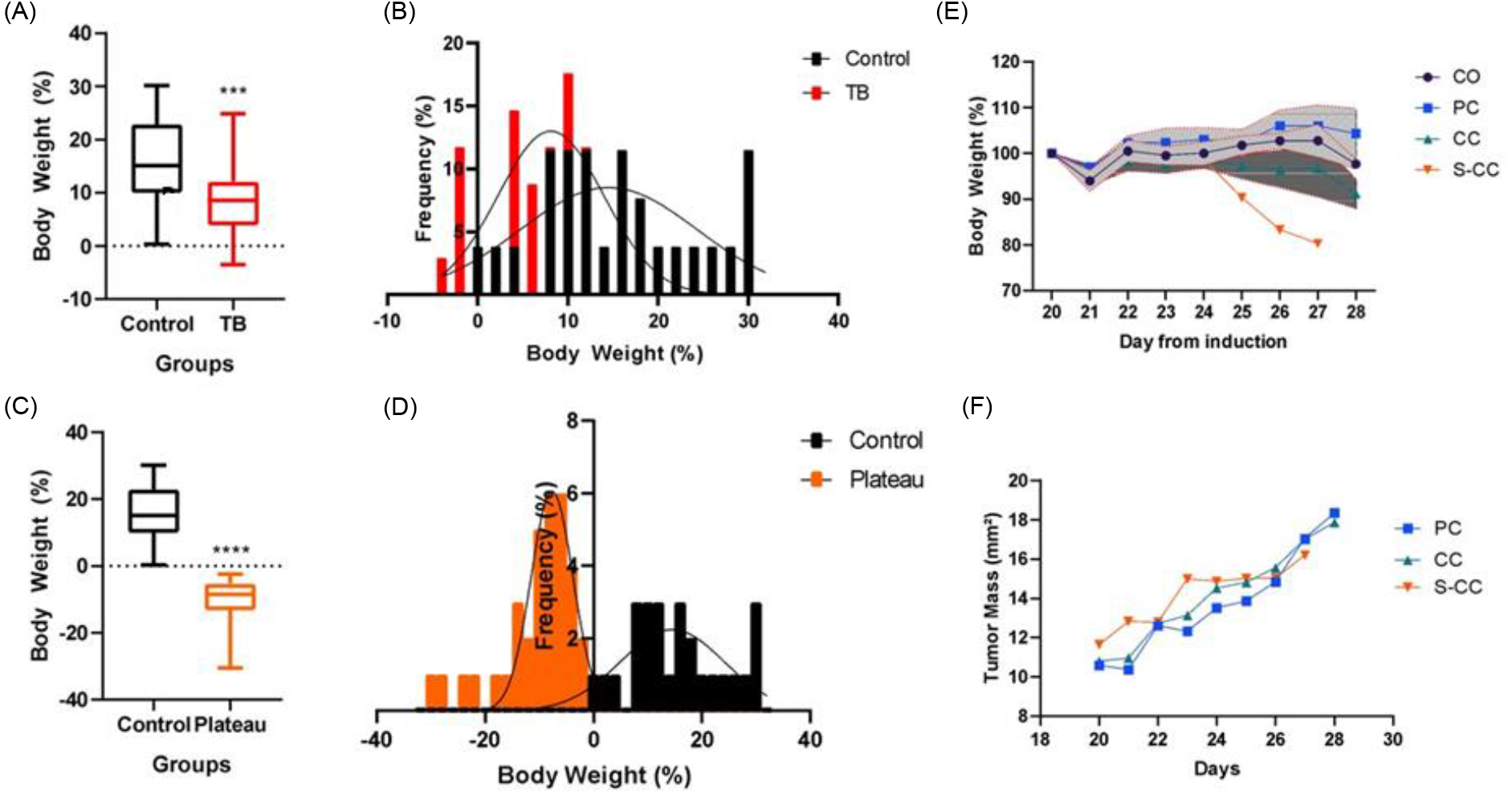

**Figure 6.**
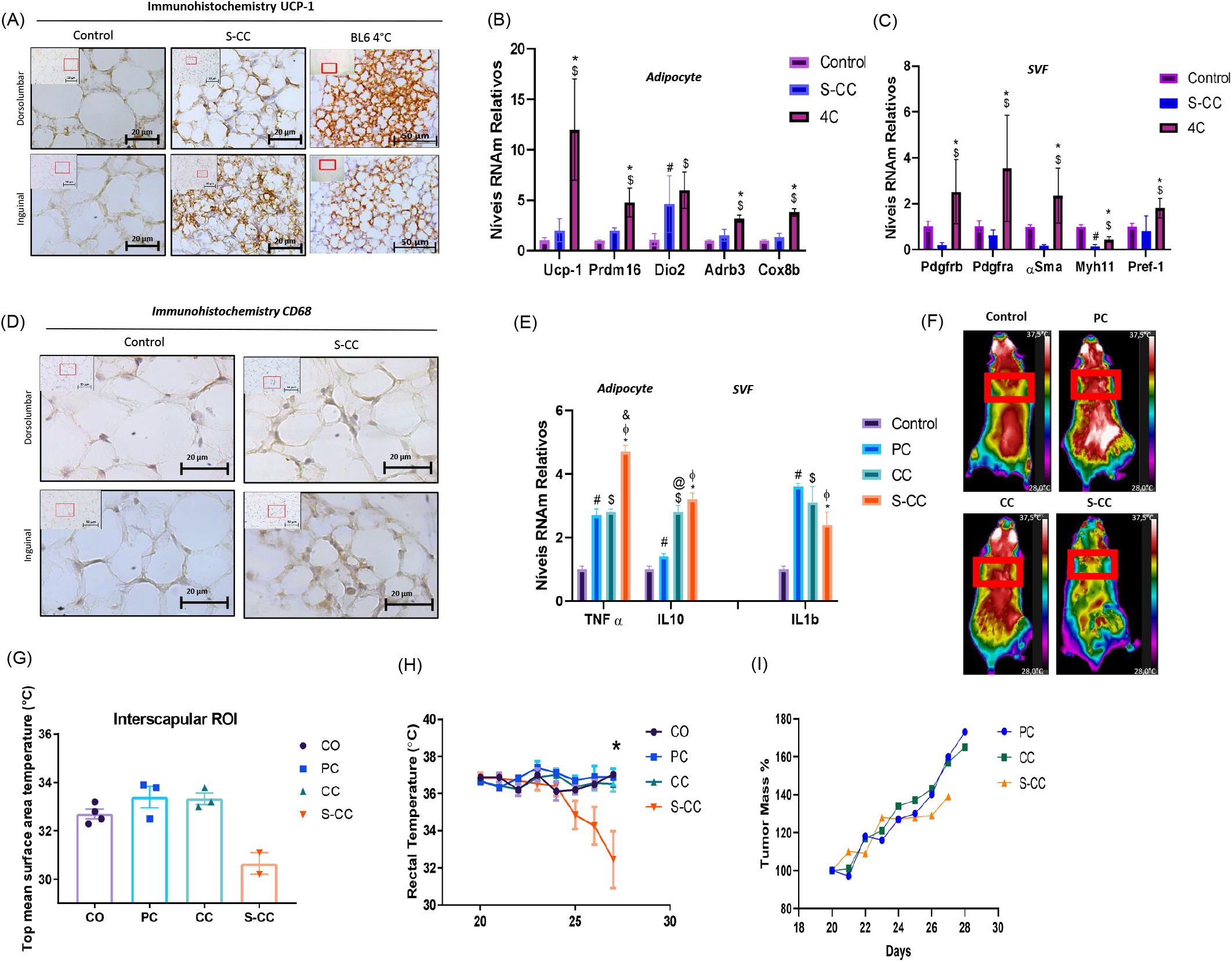

### Discussion

This study presents a syngeneic Lewis LLC1 mouse model that demonstrates distinct sequential stages of cancer cachexia development. Accessing a large cohort allowed us to identify that tumor-bearing mice do not develop the syndrome homogeneously. Based on the mice weight loss grading system, the animals can be classified according to the magnitude of their primary symptoms. This system is based on the staged cachexia classification system used in humans [23]. As a result, BW loss could be detected more accurately, and animals could be separated based on the degree of body wasting. Thus, although the classic cachexia phenotype already exists in CC animals, it is consolidated only in S-CC animals. Alterations associated with WAT remodeling have already been observed in precachectic animals (PC), which do not present alterations in BW induced by LLC1 at this astage. Moreover, using novel adipocyte culture techniques and different in vitro tumor models, it was possible to characterize cachexia-induced beige cells, suggesting that they may be the result of adipocyte transdifferentiation.

CAC, despite being described more than 100 years ago, has only recently systematically used a classification system according to the degrees of dysfunction generated by cachexia [23]. However, this system is predominantly used in clinical studies [22]. Taking advantage of the concept of grading the syndrome in cancer patients [23, 37], we adapted it for mice using three cachexia stage, according to the magnitude of body weight loss, and normalized it according to growth plateau of the tumor-bearing animal. Initially, BW loss is used as a primary criterion for classifying experimental groups. In particular, the adoption of this system results in a more homogeneous group of animals concerning the manifestation of the syndrome and separating them from those that do not develop the syndrome or develop it in an exacerbated manner. Despite the advantages of the classification system presented here, we are aware of the limitations of our study. Mainly because the experimental design needs a high number of animals. Moreover, considering the humanization criteria adopted, few animals from the S-CC group reach the end of the experiment since they must be sacrificed before the experimental period. In general, considering the established endpoints, S-CC animals are those that presented a cachexia phenotype similar to the stage of cachexia in humans.

WAT remodeling in cachexia is generally characterized by *pathologic* morphofunctional changes which take place in the early stages of the syndrome and progress to severe dysfunction [10, 38]. In our study, WAT was observed to undergo the CAC syndrome at an early stage. Several previous studies have confirmed this finding [8-10, 12]. Nevertheless, what is new is the discovery of initial signals of WAT remodeling in PC animals. Only scAT displayed a loss in its tissue mass in CC, followed by a positive correlation with BW loss. That fact was not observed in other visceral tissues. To obtain deeper information about the morphofunctional modifications in the scAT from CC group, we analyzed morphological changes and dorsal and inguinal anatomical areas. Previous study [27] has used this approach and observed that there is a prominent variation of adipose cell biogenesis when animals are exposed to the chronic cold protocol, showing that there is an increase in UCP-1 positive cells in the inguinal region. Using this approach, it was possible to observe an adipocyte atrophy in PC animals mainly in the inguinal region when compared to their respective control (non-tumor-bearing mice). Additionally, an increase in total collagen content only in the inguinal region. Furthermore, the set of results demonstrate a continuing WAT remodeling induced by cachexia; increased collagen fiber content, and fibronectin expression, well demonstrated in S-CC. PC animals also displayed high levels of main lipolytic genes followed by decrease of genes related to lipogenesis, indicating a shift the balance in lipid turnover towards to impaired uptake [10, 39]. It is important to point out that PC animals that have not yet shown BW loss or scAT atrophy, already show morphofunctional alterations before the establishment of the classic signs of cachexia. These results are in line with results presented in cancer patients without cachexia, regarding the increase in genes related to the lipolytic pathway and ECM remodeling [40].

It has been demonstrated in various experimental models that beige remodeling is an important component of accelerated energy expenditure and weight loss in mice models of CAC [1, 8, 9]. It is important to note, however, that some other studies did not find evidence of beige remodeling. In most cases, it is related to the experimental model chosen, the number of tumor cells initially injected (CAC model), and the anatomical region of WAT being examined [1, 7, 41]. In our study, we demonstrated that the anatomical location of the WAT and the severity of cachexia affect the detection of beige cells based on the mice weight loss grading system. Multilocular adipocytes were detected in scAT only in S-CC, followed by UCP-1 positive cells. Furthermore, this profile was more prominent in the inguinal region than in the dorsal-lumbar region. Although there are no preclinical studies based on our CAC grading approach, it has consistently been demonstrated that the browning phenotype exists near the endpoint (time-course studies) [2, 8], as well as in multi-experimental CAC models [9]. Interestingly, isolated adipocytes from the three experimental groups (PC, CC, and S-CC) showed increased levels of markers such as *Dio2, Prdm16*, and *Pgc1a*. In this regard, CAC will likely exhibit “classical” beige cells, i.e., thermogenesis that is mediated by UCP-1. It is interesting to note that, despite the increase in WAT browning in S-CC animals, the reduction in rectal temperature in this animal induced by CAC could be interpreted as a defense mechanism against lower body temperatures. A similar finding was reported by Petruzzeli et al. (2014), who found that WAT browning in cachectic mice is not influenced by external thermal stimuli.

Furthermore, we examined the stromal vascular fraction (SVF) cells from scAT. Markers of progenitor cells, such as *Pdgfrb, Asma*, and *Myh11*, were downregulated in the CAC, while markers of committed preadipocytes, such as *Pref-1*, were not. Genes related to adipogenesis were also evaluated for information about progenitor cells compromised by adipogenesis, which were also reduced. The presence of beige cells *(Cd137*) committed to thermogenic programming increased levels, however, was only found in S-CC. Cells that commit to “white” adipogenesis and mural progenitor cells, as well as cells that do not commit to “white” adipogenesis, showed mRNA reduced levels, suggesting that adipogenesis has deteriorated. These results are already detected in the CP group, suggesting that early changes in scAT are at the very beginning of the CAC. This study confirms previous clinical research (Fouladium)[5], in vivo [10], in vitro [40] analysis.

Various adipocyte culture models have been developed over the last several decades in order to mimic the function and morphology of adipocytes in fat depots in vivo [42]. These methods include two-dimensional (2D), freshly isolated mature fat cells, and three-dimensional culture methods. Moreover, we sought to analyze, with enhanced precision, the various aspects associated with the characterization of beige cells. Having demonstrated previously that LLC1 cells induce beige adipogenesis in a co-culture system [43], our next step was to determine whether LLC1 cell secretomes are exclusively required. In order to explore the specificity of LLC1 cells, we added conditioned media containing C-26 and MC38 tumor cells. The first relates to the induction of cachexia, which is often associated with a lack of beige cells [41, 44] and the second relates to the non-induction of CAC [41] in mice. Both tumor cells secreted factors that impaired adipogenesis in SVF primary cells in vitro. Meanwhile, only LLC1 secretome was able to increase the levels of thermogenic programming genes, including *Ucp-1, Pgc1a, Cptb*, and *Cox8b*. Even though this result has already been demonstrated [8], our experimental design proved to be successful in demonstrating two novelties: 1) the specificity of this effect in relation to LLC1 cells, and 2) the adipogenic programming is induced even without the addition of differentiation-inducing cocktail (adipogenesis). As a result, it is hypothesized that the LLC1 secretome contains components that promote beige adipogenesis over white adipogenesis. However, further studies are required both to characterize the components of these factors as well as to determine the cell lines that are sensitive to them.

Although WAT beige remodeling has been observed in different experimental models, suggesting that cachexia is associated with beige adipocytes [33, 45, 46], the origin of beige adipocytes remains a mystery. Furthermore, the majority of analyses have focused on vivo morphological characterization, as well as different in vitro assays using differentiated adipocytes for the detection of beige adipocytes and de novo adipogenesis. Aware of this scenario, we hypothesized whether beige cells arise through reprogramming or trans-differentiation of mature white adipocytes in CAC. The LLC1 secretome increased the number of UCP1-GFP+ cells in both, ex vivo and isolated mature adipocytes. It depends on how much (concentration) conditioned medium (LLC-CM) is used. Due to S-CC mice having the highest levels of WAT browning, this approach was used. Our study was conducted using the approach that allows for the cultivation of freshly mature adipocytes (MAACs) in cells derived from UCP-Cre mice [36]. Therefore, we had characterized three (3) new facts; 1-the transdifferentiation from white to beige adipocytes appears to be a relevant event in the induction of browning in cachexia, even in vitro, and 2, this fact appears to be related to the “magnitude” or intensity of cachexia, and, 3, this fact is restricted to the LLC1 secretome.

## Conclusion

Using a humanized CAC classification system, it was possible to characterize the onset of the syndrome more homogeneously in accordance with the presence of the main CAC symptoms. Earlier data showing early WAT remodeling was corroborated by this approach. Furthermore, it has been shown that scAT is affected prior to visceral depots, highlighting the importance of this depot in the development of the syndrome. Pathological beige remodeling occurred early in the course of the disease and exhibited phenotypic characteristics specific to the secretomes of LLC1 cells, depending on the severity of the disease and the anatomical location of the scAT. This scenario involves mainly the trans-differentiation or reprogramming of mature white adipocytes to become beige cells. Based on these findings, it appears that inhibiting WAT browning might be a promising approach to alleviating the severity of cachexia in cancer patients.

## Acknowledgements

We thank all members of the Laboratory of Adipose Tissue Biology for helpful discussions and critical reading of the manuscript. This work was supported by São Paulo Research Foundation (FAPESP) Grants: 2010/51078-1, 2015/19259-0 and CNPq 311966/2015-2 to M.L.B.Jr and from This work was supported by grants from NIH (DK117161 and DK117163 to SRF, and P30-DK-046200 to Adipose Biology and Nutrient Metabolism Core of Boston Nutrition and Obesity Research Center) by the Coordination for the Improvement of Higher Education Personnel, CAPES (scholarship 157207/2014-5 to K.B.N.H.S). The contents of this article are solely the responsibility of the authors and do not necessarily represent the official views of FAPESP.

## Supplementary Methods

### Tumor and Tissues collection

Lung Lewis carcinoma (LLC1, CRL-1642 -ATCC) cells were used for inducing cancer cachexia syndrome. LLC1 was injected subcutaneously into the right flanks (200 μl LLC cells 3.5x10^5^). Non-tumor bearing control mice received Phosphate-buffered saline (PBS) only. Development of CAC was monitored by tumor size and body weight for 28 days. The impact of tumor growth and development of cachexia on animal welfare were considered, with appropriate measures to monitor and alleviate suffering implementation according to adopted endpoint criteria [2]. Overnight-fasted mice were euthanized by decapitation without anesthesia. After euthanasia, subcutaneous adipose tissue (scAT) was carefully dissected and weighed. All the mice were checked if they are positive for the presence of metastasis in the lungs, as well as other visceral organs. Mesenteric adipose tissue (meAT), after careful removal of adjacent lymph nodes) and retroperitoneal adipose tissue (rpAT) were removed, weighed, snap-frozen in liquid nitrogen. If there is a presence of metastasis, the animals are excluded from the experimental group (end point exclusion criteria).

### Primary white adipocyte culture

Inguinal stroma-vascular fractions were obtained from 30–35-day-oldmalemice using the following procedure. Inguinal fat tissue was dissected, washed with PBS, minced, and digested for 45min at 37 °C in PBS containing 10mMCaCl_2_, 2.4 Uml^-1^ dispase II (Roche) and 1.5 Uml^-1^ collagenase D (Roche). Digested tissue was filtered through a 100-mm cell strainer and centrifuged at 600g for 5 min to pellet the stroma-vascular cells. The pellets were then resuspended in adipocyte culture medium (DMEM/F12 (1:1, Invitrogen) plus GlutaMAX, penicillin and streptomycin, and 10% FBS), filtered through a 40-mm cell strainer, centrifuged as above, resuspended in adipocyte culture medium and plated. The stroma-vascular cells were grown to confluence for adipocyte differentiation, which was induced by an adipogenic cocktail containing 1uM dexamethasone, 5ug.ml^-1^ insulin, 0.5uM isobutylmethylxanthine (DMI) and 1uM rosiglitazone in adipocyte culture medium. Two days after induction, cells were maintained in adipocyte culture medium containing 5 ug.ml^-1^ insulin and 1 uM rosiglitazone. Starting on day 0, cells were maintained in adipocyte culture medium only and treated with conditioned medium from Lung Lewis carcinoma (LLC1, CRL-1642 -ATCC), colorectal carcinoma 26 (C26, CT26.WT -CRL-2638 -ATCC) and colon adenocarcinoma (MC-38, CVCL_B288) tumor cells.

### Cold exposure

The animals were randomly separated into groups with at least 4 animals in each cage. Before exposure to 4°C, mice were kept at 18°C for 1 week for adaptation, since directly transferring mice from room temperature (usually 22°C) to 4°C can lead to a high mortality rate. After adaptation, the prolonged cold exposure protocol was performed (LIM et al., 2012). In the chronic cold exposure trial, the animals remained for 7 days in the cold chamber (4°C), and the temperature was checked at the beginning and end of the experimental period. A rectal thermometer (ATP® thermometer DT-610B) was used to measure body temperature.

### Histological analysis

All steps of the histological protocol were performed manually. The scAT was divided into 3 different regions (superior, middle and inferior). scAT fragments were fixed in 1 mL of HistChoice (pH 7.4), in 2 mL capacity microtubes, for 12 hours. Subsequently, the samples were submitted to the dehydration process, carried out by immersion in ethanol alcohol with increasing concentrations, 70%, 95% and 100%, for 30 minutes, and xylene for 2 hours. Then, the samples were immersed in Paraplast X-tra paraffin (Polyscience, Illinois, USA) overnight at 56ºC and embedded in Tissue Cassettes (ThermoFisher Scientific). The samples were cut on a rotary microtome (Leica RM2235, Aotec Scientific Instruments Ltda, São Paulo, SP), 5 μm thick, and positioned on slides. The sections were deparaffinized, with two successive 30-minute xylene baths, followed by hydration in ethyl alcohol baths (100%, 95%, and 70%, respectively), emerged for 5 minutes, and finally washed in running distilled water for 1 minute. The slides were stained with hematoxylin (3 minutes) and eosin (5 seconds) and then sealed with Entellan® (Merck Millipore Corporation, Massachusetts, USA) and viewed under a Leica DM 750 microscope (Aotec Scientific Instruments).

### Image analysis

The analysis of adipocyte morphology in the histological slides was performed by capturing digitized images, which were obtained using a Leica DM750 optical microscope with DFC 450 camera. For the perimeter and sectional area, 100 adipocytes were measured per section, with 5 images for each animal. To perform the morphometric measurements, Adiposoft (ImageJ) was used, and its average frequency of adipocyte diameter was 200 μm.

### Picrosirius red staining

The staining with Picrosirius Red, as a marker of total collagen, was performed immediately after the process of slide preparation. The previously fixed tissues were stained for one hour with Picrosirius Red. After staining, the plates were closed with Entellan® (Merck Millipore Corporation, Massachusetts, USA) and viewed under a Leica DM 750 microscope (Aotec Scientific Instruments). Quantification was performed using Image J software.

### Immunohistochemistry

The fragments were collected, fixed and dehydrated, as previously mentioned. For the immunohistochemistry reaction, the Histofine One Detection System: HRP Polymer (Thermo Scientific®) was used, always performing their respective positive and negative controls. The antigenic recovery was done by inducing heat for 50 minutes at 93ºC in all slides produced. Subsequently, endogenous peroxidase was blocked (hydrogen peroxide 3% for 15 minutes), and protein block (Serum Anti-Goat -KPL). The specific primary antibody (anti-UCP-1, Abcam 10983 and CD68 Abcam 125212) was used, in overnight incubation at 4°C. After incubation, Histofine One Detection polymer was applied for 20 minutes. The slides were revealed with the chromogen diaminobenzidine (DAB), and counterstained with hematoxylin. Finally, the slides were dehydrated and mounted in Entellan® synthetic medium (Merck Millipore Corporation, Massachusetts, United States of America).

### RNA isolation and qRT-PCR

Total RNA was isolated from scAT using QIAzol Lysis Reagent Protocol (QIAGEN) following the manufacturer’s instructions. cDNA was synthesized from 2 μg of total RNA using iScript cDNA Synthesis Kit (BioRad). RPL19 served as control for internal reference gene. Primer sequences used for qRT-PCR analyses were listed in Table. Analyses of qRT-PCR products were performed with the Prism 7500 SDS software (Thermo Fisher Scientific). Relative quantification of mRNA amount was obtained by the by 2^−(ΔΔCt)^ method.

### Primer selection

The primer pairs, used in the qPCR technique, were synthesized by Invitrogen -Life Technologies and Síntese Biotecnologia. The sequences of the primers were determined using the information of the target gene, deposited in the GenBank platform (https://www.ncbi.nlm.nih.gov/nucleotide/). From the information contained in GenBank, the DNA sequence of the target gene, as well as the mRNA sequence, were obtained. Thus, all primer pairs suggested by the software were compared, and their specificity was determined on the Nucleotide Blast platform (available at: https://blast.ncbi.nlm.nih.gov/). Only primers that achieved high specificity with the target gene were selected.

### Heat dissipation measurement (thermography)

Thermographic assay was used to measure the approximate temperature of the BAT. The protocol was adapted from Crane and collaborators [19], and a FlirT 460 camera was used to obtain the thermographic images. For the standardization of the room, an acclimatization was performed for 10 minutes, in which the animals were placed in the image acquisition room. After completion of the acclimatization, images were acquired individually, i.e., one animal at a time. The mouse was positioned in the previously stipulated demarcation (60 cm away from the camera). Two thermography sessions were performed on the first and last day of the experiment. The images were analyzed using Flir Tools software. A minimum (24°C) and maximum (37.5°C) temperature standard was established, and subsequently, the regions of interest were established, the first referring to the BAT, called scapular region of interest (ROI), and the body region of interest (ROI), which was established below the ROI, up to the base of the tail.

## References

1. Henriques F, Júnior MLB. Adipose Tissue Remodeling during Cancer-Associated Cachexia: Translational Features from Adipose Tissue Dysfunction %J Immunometabolism. 2020;2:e200032. doi:10.20900/immunometab20200032

2. Henriques F, Lopes MA, Franco FO, Knobl P, Santos KB, Bueno LL, et al. Toll-Like Receptor-4 Disruption Suppresses Adipose Tissue Remodeling and Increases Survival in Cancer Cachexia Syndrome. Sci Rep. 2018;8:18024. doi:10.1038/s41598-018-36626-3

3. Anderson LJ, Lee J, Anderson B, Lee B, Migula D, Sauer A, et al. Whole-body and adipose tissue metabolic phenotype in cancer patients. J Cachexia Sarcopenia Muscle. 2022;13:1124–33. doi:10.1002/jcsm.12918

4. Tsoli M, Moore M, Burg D, Painter A, Taylor R, Lockie SH, et al. Activation of thermogenesis in brown adipose tissue and dysregulated lipid metabolism associated with cancer cachexia in mice. Cancer Res. 2012;72:4372–82. doi:10.1158/0008-5472.CAN-11-3536

5. Fouladiun M, Korner U, Bosaeus I, Daneryd P, Hyltander A, Lundholm KG. Body composition and time course changes in regional distribution of fat and lean tissue in unselected cancer patients on palliative care--correlations with food intake, metabolism, exercise capacity, and hormones. Cancer. 2005;103:2189–98. doi:10.1002/cncr.21013

6. Batista ML, Jr., Henriques FS, Neves RX, Olivan MR, Matos-Neto EM, Alcantara PS, et al. Cachexia-associated adipose tissue morphological rearrangement in gastrointestinal cancer patients. J Cachexia Sarcopenia Muscle. 2016;7:37–47. doi:10.1002/jcsm.12037

7. Bing C, Russell S, Becket E, Pope M, Tisdale MJ, Trayhurn P, et al. Adipose atrophy in cancer cachexia: morphologic and molecular analysis of adipose tissue in tumour-bearing mice. Br J Cancer. 2006;95:1028–37. doi:10.1038/sj.bjc.6603360

8. Kir S, White JP, Kleiner S, Kazak L, Cohen P, Baracos VE, et al. Tumour-derived PTH-related protein triggers adipose tissue browning and cancer cachexia. Nature. 2014;513:100–4. doi:10.1038/nature13528

9. Petruzzelli M, Schweiger M, Schreiber R, Campos-Olivas R, Tsoli M, Allen J, et al. A switch from white to brown fat increases energy expenditure in cancer-associated cachexia. Cell Metab. 2014;20:433–47. doi:10.1016/j.cmet.2014.06.011

10. Henriques FS, Sertie RAL, Franco FO, Knobl P, Neves RX, Andreotti S, et al. Early suppression of adipocyte lipid turnover induces immunometabolic modulation in cancer cachexia syndrome. FASEB J. 2017;31:1976–86. doi:10.1096/fj.201601151R

11. Batista ML, Jr., Neves RX, Peres SB, Yamashita AS, Shida CS, Farmer SR, et al. Heterogeneous time-dependent response of adipose tissue during the development of cancer cachexia. J Endocrinol. 2012;215:363–73. doi:10.1530/JOE-12-0307

12. Franco FO, Lopes MA, Henriques FS, Neves RX, Bianchi Filho C, Batista ML, Jr. Cancer cachexia differentially regulates visceral adipose tissue turnover. J Endocrinol. 2017;232:493–500. doi:10.1530/JOE-16-0305

13. Petruzzelli M, Wagner EF. Mechanisms of metabolic dysfunction in cancer-associated cachexia. Genes Dev. 2016;30:489–501. doi:10.1101/gad.276733.115

14. Betancourt A, Busquets S, Ponce M, Toledo M, Guardia-Olmos J, Pero-Cebollero M, et al. The animal cachexia score (ACASCO). Animal Model Exp Med. 2019;2:201–9. doi:10.1002/ame2.12082

15. Wallace J. Humane endpoints and cancer research. ILAR J. 2000;41:87–93. doi:10.1093/ilar.41.2.87

16. Rodbell M. Metabolism of Isolated Fat Cells. I. Effects of Hormones on Glucose Metabolism and Lipolysis. J Biol Chem. 1964;239:375–80.

17. Henriques FdS. CARACTERIZAÇÃO DO PAPEL DO TLR4 NO REMODELAMENTO DO TECIDO ADIPOSO DURANTE O DESENVOLVIMENTO DA CAQUEXIA ASSOCIADA AO CÂNCER. 2016;115.

18. Voltarelli FA, Frajacomo FT, Padilha CS, Testa MTJ, Cella PS, Ribeiro DF, et al. Syngeneic B16F10 Melanoma Causes Cachexia and Impaired Skeletal Muscle Strength and Locomotor Activity in Mice. Front Physiol. 2017;8:715. doi:10.3389/fphys.2017.00715

19. Crane JD, Mottillo EP, Farncombe TH, Morrison KM, Steinberg GR. A standardized infrared imaging technique that specifically detects UCP1-mediated thermogenesis in vivo. Mol Metab. 2014;3:490–4. doi:10.1016/j.molmet.2014.04.007

20. Ray MA, Johnston NA, Verhulst S, Trammell RA, Toth LA. Identification of markers for imminent death in mice used in longevity and aging research. J Am Assoc Lab Anim Sci. 2010;49:282–8.

21. Junqueira LC, Toledo OM, Montes GS. Histochemical and morphological studies on a new type of acellular cartilage. Basic Appl Histochem. 1983;27:1–8.

22. Baracos VE, Martin L, Korc M, Guttridge DC, Fearon KCH. Cancer-associated cachexia. Nat Rev Dis Primers. 2018;4:17105. doi:10.1038/nrdp.2017.105

23. Blum D, Stene GB, Solheim TS, Fayers P, Hjermstad MJ, Baracos VE, et al. Validation of the Consensus-Definition for Cancer Cachexia and evaluation of a classification model--a study based on data from an international multicentre project (EPCRC-CSA). Ann Oncol. 2014;25:1635–42. doi:10.1093/annonc/mdu086

24. Fearon K, Arends J, Baracos V. Understanding the mechanisms and treatment options in cancer cachexia. Nat Rev Clin Oncol. 2013;10:90–9. doi:10.1038/nrclinonc.2012.209

25. Fearon K, Strasser F, Anker SD, Bosaeus I, Bruera E, Fainsinger RL, et al. Definition and classification of cancer cachexia: an international consensus. Lancet Oncol. 2011;12:489–95. doi:10.1016/S1470-2045(10)70218-7

26. Batista ML, Jr., Peres SB, McDonald ME, Alcantara PS, Olivan M, Otoch JP, et al. Adipose tissue inflammation and cancer cachexia: possible role of nuclear transcription factors. Cytokine. 2012;57:9–16. doi:10.1016/j.cyto.2011.10.008

27. Chi J, Wu Z, Choi CHJ, Nguyen L, Tegegne S, Ackerman SE, et al. Three-Dimensional Adipose Tissue Imaging Reveals Regional Variation in Beige Fat Biogenesis and PRDM16-Dependent Sympathetic Neurite Density. Cell Metab. 2018;27:226–36 e3. doi:10.1016/j.cmet.2017.12.011

28. Cowin SC. Tissue growth and remodeling. Annu Rev Biomed Eng. 2004;6:77–107. doi:10.1146/annurev.bioeng.6.040803.140250

29. Itoh M, Suganami T, Hachiya R, Ogawa Y. Adipose tissue remodeling as homeostatic inflammation. Int J Inflam. 2011;2011:720926. doi:10.4061/2011/720926

30. Bonet ML, Oliver P, Palou A. Pharmacological and nutritional agents promoting browning of white adipose tissue. Biochim Biophys Acta. 2013;1831:969–85. doi:10.1016/j.bbalip.2012.12.002

31. Jiang Y, Berry DC, Graff JM. Distinct cellular and molecular mechanisms for beta3 adrenergic receptor-induced beige adipocyte formation. Elife. 2017;6: doi:10.7554/eLife.30329

32. Sidossis L, Kajimura S. Brown and beige fat in humans: thermogenic adipocytes that control energy and glucose homeostasis. J Clin Invest. 2015;125:478–86. doi:10.1172/JCI78362

33. Wu J, Bostrom P, Sparks LM, Ye L, Choi JH, Giang AH, et al. Beige adipocytes are a distinct type of thermogenic fat cell in mouse and human. Cell. 2012;150:366–76. doi:10.1016/j.cell.2012.05.016

34. Himms-Hagen J, Melnyk A, Zingaretti MC, Ceresi E, Barbatelli G, Cinti S. Multilocular fat cells in WAT of CL-316243-treated rats derive directly from white adipocytes. Am J Physiol Cell Physiol. 2000;279:C670–81. doi:10.1152/ajpcell.2000.279.3.C670

35. Tisdale MJ. Catabolic mediators of cancer cachexia. Curr Opin Support Palliat Care. 2008;2:256–61. doi:10.1097/spc.0b013e328319d7fa

36. Harms MJ, Li Q, Lee S, Zhang C, Kull B, Hallen S, et al. Mature Human White Adipocytes Cultured under Membranes Maintain Identity, Function, and Can Transdifferentiate into Brown-like Adipocytes. Cell Rep. 2019;27:213–25 e5. doi:10.1016/j.celrep.2019.03.026

37. Fearon KC, Voss AC, Hustead DS, Cancer Cachexia Study G. Definition of cancer cachexia: effect of weight loss, reduced food intake, and systemic inflammation on functional status and prognosis. Am J Clin Nutr. 2006;83:1345–50. doi:10.1093/ajcn/83.6.1345

38. Arner P. Medicine. Lipases in cachexia. Science. 2011;333:163–4. doi:10.1126/science.1209418

39. Arner P, Langin D. Lipolysis in lipid turnover, cancer cachexia, and obesity-induced insulin resistance. Trends Endocrinol Metab. 2014;25:255–62. doi:10.1016/j.tem.2014.03.002

40. Alves MJ, Figueredo RG, Azevedo FF, Cavallaro DA, Neto NI, Lima JD, et al. Adipose tissue fibrosis in human cancer cachexia: the role of TGFbeta pathway. BMC Cancer. 2017;17:190. doi:10.1186/s12885-017-3178-8

41. Wu L, Zhang L, Li B, Jiang H, Duan Y, Xie Z, et al. AMP-Activated Protein Kinase (AMPK) Regulates Energy Metabolism through Modulating Thermogenesis in Adipose Tissue. Front Physiol. 2018;9:122. doi:10.3389/fphys.2018.00122

42. Dufau J, Shen JX, Couchet M, De Castro Barbosa T, Mejhert N, Massier L, et al. In vitro and ex vivo models of adipocytes. Am J Physiol Cell Physiol. 2021;320:C822–C41. doi:10.1152/ajpcell.00519.2020

43. Lopes MA, Oliveira Franco F, Henriques F, Peres SB, Batista ML, Jr. LLC tumor cells-derivated factors reduces adipogenesis in co-culture system. Heliyon. 2018;4:e00708. doi:10.1016/j.heliyon.2018.e00708

44. Bing C. Lipid mobilization in cachexia: mechanisms and mediators. Curr Opin Support Palliat Care. 2011;5:356–60. doi:10.1097/SPC.0b013e32834bde0e

45. Giralt M, Villarroya F. White, brown, beige/brite: different adipose cells for different functions? Endocrinology. 2013;154:2992–3000. doi:10.1210/en.2013-1403

46. Harms M, Seale P. Brown and beige fat: development, function and therapeutic potential. Nat Med. 2013;19:1252–63. doi:10.1038/nm.3361

